# Estimating the frequency of multiplets in single-cell RNA sequencing from cell-mixing experiments

**DOI:** 10.1101/293639

**Authors:** Jesse D. Bloom

## Abstract

In single-cell RNA-sequencing, it is important to know the frequency at which the sequenced transcriptomes actually derive from multiple cells. A common method to estimate this multiplet frequency is to mix two different types of cells (e.g., human and mouse), and then determine how often the transcriptomes contain transcripts from both cell types. When the two cell types are mixed in equal proportion, the calculation of the multiplet frequency from the frequency of mixed transcriptomes is straightforward. But surprisingly, there are no published descriptions of how to calculate the multiplet frequency in the general case when the cell types are mixed unequally. Here I derive equations to analytically calculate the multiplet frequency from the numbers of observed pure and mixed transcriptomes when two cell types are mixed in arbitrary proportions, under the assumption that the loading of cells into droplets or wells is Poisson.

## INTRODUCTION

Many methods for single-cell RNA sequencing involve partitioning cells into barcoded droplets (Klein et al., 2015; Macosko et al., 2015; Zheng et al., 2017), wells (Gierahn et al., 2017), or combinations of wells (Cao et al., 2017). As long as the number of possible partitions exceeds the number of cells, then most partitions will contain at most one cell. However, some fraction of the non-empty partitions will contain multiple cells, and estimating this *multiplet frequency* is an important aspect of experimental quality control.

The most common method to determine the multiplet frequency is to use a mix two types of cells (e.g., human and mouse). During the analysis of the sequencing results, each non-empty partition can be identified as containing transcripts from one or both of the two cell types. Partitions that contain a substantial number of transcripts from both cell types must be multiplets. If the two cell types are mixed equally and the average number of cells per partition is low (so that most multiplets are doublets), then the multiplet frequency can be estimated as simply twice the fraction of non-empty partitions that contain a mix of cell types. The logic is that all the multiplets are doublets, and only half the doublets will have cells of both types (the others will have two cells of the same type). This approach has been used to estimate the multiplet frequency during the prototyping of most single-cell RNA sequencing methods (Klein et al., 2015; Macosko et al., 2015; Zheng et al., 2017; Gierahn et al., 2017; Cao et al., 2017).

However, in some cases the two cell types may be mixed in unequal proportions. Unequal mixing could arise simply from error during cell counting, but it can also be a desirable aspect of experimental design. For instance, if the researcher is actually interested in the human cells and simply wants to include an internal control to estimate the multiplet frequency during each new experiment, then (s)he may want to add fewer mouse cells so that most of the resulting data is for the human cells. But when the cells are mixed unequally, it is no longer valid to estimate the multiplet frequency as simply twice the fraction of non-empty partitions that contain a mix of both cell types. Surprisingly, I could find no published descriptions of how to calculate the multiplet frequency from unequal mixes of two cell types. Here I remedy this gap in the literature by deriving the equations to compute the multiplet frequency when the cells are mixed in arbitrary proportions under the assumption that the number of cells per partition is Poisson distributed.

## RESULTS AND DISCUSSION

### Derivation of multiplet frequency from observed numbers of pure and mixed-cell droplets

Consider the case in which cells of two types (e.g., human and mouse) are distributed into individual barcoded droplets, although the same logic applies if the cells are distributed into barcoded wells or combinations of wells. Assume the sequencing data have been analyzed so that each non-empty droplet can be classified as containing at least one cell of type 1, at least one cell of type 2, or cells of both types. I will refer to the number of droplets in each of these three groupings as *N*_1_, *N*_2_, and *N*_1,2_, respectively. For instance, the 10X `cellranger` pipeline (version 2.1.1) returns these numbers as the “Estimated Number of Cell Partitions.”

The only assumption of the derivation is that the number of cells per droplet is Poisson distributed. Let µ_1_ be the average number of cells of type 1 per droplet, and µ_2_ be the average number of cells of type 2 per droplet. The average number of cells of any type per droplet is then µ_1_ + µ_2_. So the probability that a droplet contains at least one cell of any type is

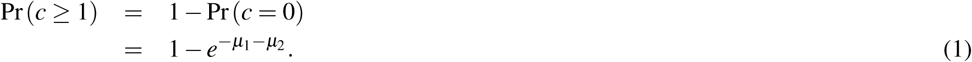

Likewise, the probability that a droplet contains multiple cells of any type (e.g., a multiplet) is

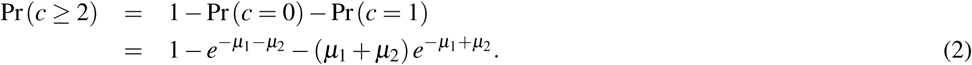

The multiplet frequency *M* is simply the probability that a droplet with at least one cell actually contains multiple cells, which is

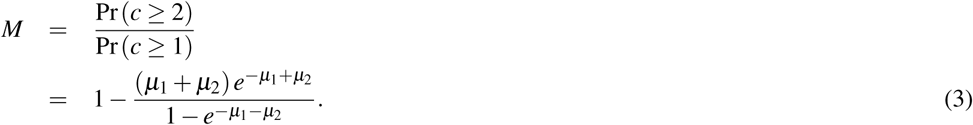

However, evaluating this expression for *M* requires the values of µ_1_ and µ_2_.

We can write down equations for µ_1_ and µ_2_ by again using the fact that the number of cells per droplet is Poisson distributed. Specifically, if *N* is the total number of droplets (empty and non-empty), then the expected number of droplets that have at least one cell of type 1 is *N* × Pr (*c*_1_ ≥ 1) = *N* (1 − *e^−µ^*^1^). The observed number of droplets with at least one cell of type 1 is *N*_1_, so setting the observed number equal to the expected number gives us an equation for µ_1_,

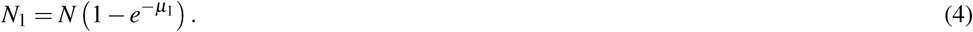

This equation is easily solved for µ_1_ to yield

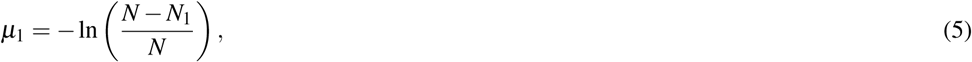

and likewise for µ_2_,

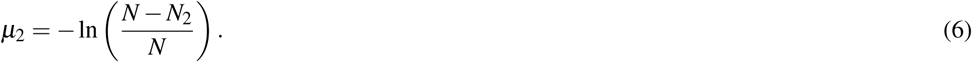

Equations 5 and 6 give us a way to determine the values (µ_1_ and µ_2_) needed to calculate the multiplet frequency (Equation 3) in terms of the experimental observables *N*_1_ and *N*_2_. Unfortunately, these two equations also require knowledge of the total (empty and non-empty) number of droplets *N*, which is not directly observable from the sequencing data.

However, we can take advantage of another relationship to calculate *N*. The fraction of all (empty and non-empty) droplets that contain cells of both types is 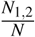, and this fraction is simply the product of the probability that a droplet contains at least one cell of type 1 with the probability that a droplet contains at least one cell of type 2, which in mathematical terms can be stated as Pr (*c*_1_ ≥ 1 ∧ *c*_2_ ≥ 1) = Pr (*c*_1_ ≥ 1) × Pr (*c*_2_ ≥ 1). Therefore,

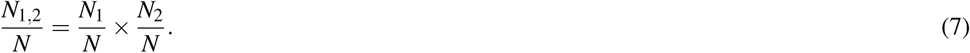

This equation can be solved to give

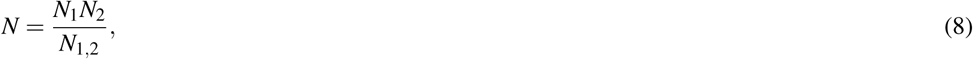

which can be completely evaluated in terms of the experimental observables. Equations 5, 6, and 8 can be used to calculate µ_1_ and µ_2_ in terms of the experimental observables, and those results used to calculate the multiplet frequency via Equation 3. This provides an analytic solution for the multiplet frequency in terms of the three experimental observables.

### Implementation and example calculations

A simple function to perform the calculations described in the previous subsection is implemented in Python and R in the Jupyter notebook found at https://github.com/jbloomlab/multiplet_freq/blob/master/calcmultiplet.ipynb (see also Supplemental files 1 and 2). To illustrate the calculations, I used this function to calculate the multiplet frequency for hypothetical data.

First, consider hypothetical data in which the two types of cells are mixed in equal proportions. Prior papers have approximated the multiplet frequency from such experiments as simply twice the fraction of non-empty droplets that contain cells of both types (Klein et al., 2015; Macosko et al., 2015; Zheng et al., 2017; Cao et al., 2017), which is 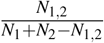 in the notation defined in the previous subsection. Table 1 shows that the exact equation derived in the previous subsection gives very similar results to this approximate method as long as the multiplet frequency is low. When the multiplet frequency becomes high, the approximate method starts to overestimate the true multiplet frequency, since it fails to account for the fact that some multiplets will contain more than two cells.

**Table 1.** Multiplet frequencies for three hypothetical experiments in which human and mouse cells are mixed equally. The multiplet frequencies calculated using the exact method described here (column *multiplet freq*) are very similar to those obtained simply by multiplying by two the fraction of non-empty droplets that contain cells of both types (column *twice cross celltype freq*). However, the two methods are slightly different at higher multiplet frequencies, since the latter method fails to account for multiplets that have more than two cells.

| experiment | human droplets | mouse droplets | nonempty droplets | human and mouse droplets | multiplet freq | twice cross celltype freq |
| --- | --- | --- | --- | --- | --- | --- |
| 1 | 2005 | 2005 | 4000 | 10 | 0.005 | 0.005 |
| 2 | 2050 | 2050 | 4000 | 100 | 0.049 | 0.050 |
| 3 | 2500 | 2500 | 4000 | 1000 | 0.425 | 0.500 |

Next, consider hypothetical data in which the two types of cells are mixed in unequal proportions. Table 2 shows the multiplet frequencies for several such experiments. An interesting aspect of the results is that at high multiplet frequencies and very unequal cell proportions, the multiplet frequency is substantially *lower* than the fraction of droplets containing the rarer cell type that contain a mix of both cell types. The reason is that multiplets (particularly higher-order ones) become more and more likely to contain at least one cell of the rarer type relative to droplets that contain only one cell. For instance, in the final experiment in Table 2, two thirds of the droplets containing mouse cells have a mix of both cell types, yet less than half the non-empty droplets are multiplets (the multiplet frequency is 0.459). This somewhat non-intuitive results illustrates the importance of using the correct mathematical relationship to calculate the multiplet frequency when cell types are mixed unequally.

**Table 2.** Multiplet frequencies for five hypothetical experiments in which human and mouse cells are mixed unequally.

| experiment | human droplets | mouse droplets | nonempty droplets | human and mouse droplets | multiplet freq |
| --- | --- | --- | --- | --- | --- |
| 1 | 2050 | 2050 | 4000 | 100 | 0.049 |
| 2 | 3050 | 1050 | 4000 | 100 | 0.065 |
| 3 | 3550 | 550 | 4000 | 100 | 0.110 |
| 4 | 3850 | 250 | 4000 | 100 | 0.245 |
| 5 | 3950 | 150 | 4000 | 100 | 0.459 |

## CONCLUSIONS

I have described how to calculate the multiplet frequency in single-cell RNA sequencing experiments in which two cell types are mixed in arbitrary proportions. It is important to note that this calculation requires that the sequencing data have already been analyzed to determine whether each partition contains a non-negligible number of transcripts from each cell type, but many common analysis programs (such as the 10X cellranger pipeline) already do this. It is also important to note that the approach in this paper only calculates the multiplet frequency—it does *not* actually identify the multiplets so that they can be removed from downstream analyses. For that purpose, other more sophisticated approaches have been developd (Ilicic et al., 2016; Stoeckius et al., 2017; Kang et al., 2018). Nonetheless, simply calculating the multiplet frequency from the data returned by standard pipelines such as the 10X cellranger is important for many purposes, and the results here enable that to be done regardless of the proportions at which the cell types are mixed.

## METHODS

The LaTex source for this paper, the Jupyter notebook that implements the calculations, and all materials associated with the writing and review of the paper are publicly available in a GitHub repository at https://github.com/jbloomlab/multiplet_freq. The Jupyter notebook is also available in Supplemental file 1, and an HTML rendering of that notebook is in Supplemental file 2.

---

**Supplemental file 1.** A Jupyter notebook that implements the calculations in Python and R functions, and does the calculations for the examples shown in the tables in this paper.

**Supplemental file 2.** This file contains an HTML rendering of the Jupyter notebook in Supplemental file 1.

